# Generation and benchmarking of a collection of hiPSC lines from Schizophrenia Patients with Diverse Clinical Profiles

**DOI:** 10.1101/2024.01.14.575590

**Authors:** Elena Rita Vecchi, Claudia Vittoria Olmeda, Daniele Bottai, Palma Finelli, Marilyn Marlene Angulo Salavarria, Cristina Gervasini, Laura Mangiaterra, Francesco Lombardi, Claudio Sanguineti, Marco Onorati, Luciano Conti, Armando D’Agostino

## Abstract

Limited therapeutic advancements in Schizophrenia (SCZ) depend on the heterogeneous nature of the disorder, impacting drug development and clinical trials that assume uniform therapy response, neglecting individual genetic and epigenomic variability. Disease modeling using human induced pluripotent stem cells (hiPSCs) is ideally suited for precision medicine, enabling individualized treatment approaches. Here, we describe the generation of patient-specific lines from somatic cells of SCZ individuals with well-defined diverse clinical trajectories using a Sendai virus-based reprogramming system. Karyotypically and CGH-array validated, the generated hiPSCs expressed diagnostic markers and demonstrated functional pluripotency. Converting these hiPSCs into neural progenitor cells enables the identification of aberrant cellular phenotypes associated with specific pathologically relevant neural phenotypes. This collection of hiPSC lines serves as a platform for developing therapeutic compounds targeting neural populations, potentially addressing early-stage disease alterations.

## INTRODUCTION

While the characteristic symptoms of Schizophrenia (SCZ) manifest between late adolescence and early adulthood in the majority of patients (Solmi et al., 2021), it is currently conceptualized as a neurodevelopmental condition, often preceded by a prodromal period that can emerge at childhood (Jauhar et al., 2022). *Postmortem* studies have revealed defects in neuronal populations, such as reduced spine density in prefrontal cortices (Dienel et al., 2022). Disruptive de novo mutations have been shown to converge in a network of co-expressed genes with a pathogenetic impact on the developing brain of those who will develop SCZ (Fromer et al., 2014). Functional genomic data strongly indicate that disruptions in synaptic connectivity, linked to susceptibility to SCZ, begin during prenatal development and persist throughout the lifespan. Defined alleles are identified as risk factors for the disorder, exerting their influence by directly impacting synaptic function in adulthood (Hall and Bray, 2022). Although research on adult *Postmortem* brain tissue and animal models provided valuable findings, the former reveal endpoints and provide limited insights into disease predisposition or initiation. On the other hand, animal studies may not fully capture the polygenic nature of the disease.

Human induced pluripotent stem cells (hiPSCs) provide a promising alternative, allowing the creation of SCZ patient-specific disease models to explore cellular and molecular phenotypes (Angulo Salavarria et al., 2023). Indeed, hiPSCs enable the generation of various brain-specific cell types, including cortical neurons, interneurons, astrocytes, and microglia (Abud et al., 2017; Muffat et al, 2016; Dell’Amico et al. 2023). Several studies identified impaired neuronal maturation, reduction of neurite length and outgrowth through these cellular populations, along with an excitatory-inhibitory imbalance, including enhanced GABAergic-related neuronal expression and genetic upregulation (Brennand et al., 2011; Park et al., 2020; Sawada et al., 2020). Other studies reported mitochondrial functional impairments, irregular cellular distribution of organelles, dissipation of mitochondrial membrane potential, reduced ATP levels, and disruptions in mitochondrial network structure and connectivity (for a comprehensive review, see Dubonyte et al., 2022).

Of note, this novel hiPSC-based disease modelling may prove to be particularly suited for precision approaches to individually tailored treatments. It has become increasingly clear that overall clinical effectiveness of available treatments is limited by the highly heterogeneous nature of SCZ. The variability of clinical phenotypes, course and treatment response has a significant impact on both drug development and clinical trials, which are inherently flawed by the assumption that all individuals should respond uniformly to a given therapy, overlooking the intricate genetic and epigenomic variations that shape patient individuality (Spark et al., 2022).

This study reports the generation of a collection of validated hiPSC lines from SCZ patients with distinct clinical phenotypes, including course and treatment response profiles. The generation of patient-derived hiPSC will consent to investigate the molecular-dependent mechanisms behind the genetic background and so the development of new personalized treatments.

## METHODS AND MATERIALS

### Study population

Blood samples were collected from a cohort of patients who participated in the SPINDLE-2 study at the Department of Mental Health and Addiction of the San Paolo University Hospital in Milan, Italy. The study was approved by the Milan Area 1 Ethics Committee (Prot. N° 34420/2019) and all patients provided informed consent. All study procedures were conducted according to the Declaration of Helsinki.

P28LR (Treatment–Resistant Schizophrenia with excellent response to clozapine): Caucasian male diagnosed with Treatment-Resistant Schizophrenia (TRS), 27 year-old at the time of sampling. First hospitalization at the onset of psychosis, when he was 23 year-old. No reported family history of psychiatric disorders. Childhood reported as uneventful, marked by commendable scholastic and social functioning, and devoid of significant medical conditions.

Two major stressful events reported at the age of 16 (parents’ divorce and a change of school). Concurrently, he began occasional cannabis use. Initial treatment with aripiprazole and low-dose haloperidol yielded partial response and was discontinued by the patient. The patient eventually had an excellent and sustained response to low-dose clozapine (150 mg), allowing him to pursue university studies, work as a chess teacher, and maintain robust social functioning (Global Assessment of Functioning, GAF = 80%).

P25FB (Early-Onset Schizophrenia with poor treatment response and unfavourable course): North African female who relocated to Italy during childhood, diagnosed with EOS, 21 year-old at the time of sampling. Positive family history for non–affective psychosis (mother). Not known medical or physical conditions and a brief history of cannabis abuse concurrent with psychosis onset. From childhood, she displayed a tendency to avoid social contact and experienced difficulties in forming relationships with peers. Psychosis onset resulting in her first hospitalization at the age of 17 was preceded by approximately one year of subthreshold symptoms following a stressful life event (death of an uncle with whom she had an elective affective bond). Over the years, the patient failed to respond to various antipsychotic drugs (olanzapine, aripiprazole, paliperidone, lurasidone). At the time of sampling, the patient was still unemployed and exhibiting poor social functioning (GAF = 60%).

P21EV (Schizophrenia, responder to long–acting injectable aripiprazole): South–East Asian woman residing in Italy, 44 year-old at the time of sampling. Born with no reported medical conditions or family history of psychiatric disorders, she was first hospitalized at the age of 39 due to acute psychosis. Over the previous three years, the patient displayed worsening signs of social withdrawal, grossly disorganized behavior, and neglect of self and surroundings following an intensely stressful life event (divorce and loss of custody of her children). Disorganized behavior had likely commenced in the perinatal period and auditory hallucinations surfaced five months postpartum, with concurrent methamphetamine use. Initially treated with paliperidone, which was switched to aripiprazole due to invalidating side effects such as akathisia, hyperprolactinemia, and emotional blunting. Aripiprazole was administered orally (30 mg daily) and later through long-acting intramuscular therapy (400 mg every 28 days). The patient demonstrated sustained and favorable response to aripiprazole over subsequent years, with complete resolution of delusions and disorganization; occasionally, she experienced simple auditory hallucinations, and persistent negative symptoms (emotional blunting and avolition), and mild cognitive dysfunction; social services assisted her in securing employment as a cleaner (GAF = 61%).

P11FA (Treatment–Resistant Schizophrenia, good response to long–acting injectable aripiprazole after partial response to clozapine): North African female who relocated to Italy during childhood, 25 year-old at the time of sampling. No reported family history of major psychiatric conditions, no medical condition nor history of substance or alcohol abuse. Traumatic life events during childhood included witnessing a neighbor’s death and exposure to domestic violence. Onset of psychosis, which led to her first hospitalization, occurred when she was 21 year-old. After insufficient response to aripiprazole and risperidone (and extrapyramidal symptoms/hyperprolactinemia from the latter), she responded positively to clozapine treatment (150 mg daily). Post-discharge, the patient exhibited low insight and insufficient collaboration, leading to rehospitalization two years later after discontinuation of clozapine. After an unsuccessful trial with lurasidone, she presented adequate symptom control with aripiprazole, which was switched to long-acting formulation.

Psychiatric services assisted her in job placement as a caregiver for the elderly, and she completed a high-school diploma at the age of 25, despite a globally limited social functioning and few relationships (GAF = 71%).

P22KG (Treatment-Resistant Schizophrenia with co-morbid Internet Gaming and Alcohol Use Disorders, Clozapine non–responder): Italian–born male of Latin American descent, diagnosed with TRS, 24 year-old at the time of sampling. No reported family history of psychiatric disorders, comorbid Internet Gaming Disorder and remitted Alcohol Use Disorder. Psychosis onset leading to a first hospitalization occurred at the age of 19. The patient underwent treatment with three different antipsychotic drugs (aripiprazole, risperidone, paliperidone), discontinued due to poor adherence and/or adverse reactions. Clozapine, alone and combined with amisulpride, was discontinued due to poor adherence. Subsequently, risperidone and aripiprazole LAI formulations were administered, with risperidone LAI discontinued due to akathisia. Aripiprazole LAI resulted in satisfactory control of symptoms, but yielded marked weight gain. The patient remained unemployed and had poor social functioning (GAF 50%).

### PBMC extraction and Reprogramming to hiPSCs

Blood samples from donors with informed consent were collected into BD Vacutainer CPT cell preparation tubes and subjected to 30 min centrifugation (1800 ×g at room temperature). Peripheral Blood Mononuclear Cells (PBMCs) were purified by resuspending the resulting buffy coat in 15 mL of PBS and centrifuging it at room temperature (RT) for 15 min at 300× g. Aliquots of 2 x 10^6^ cells were cryopreserved in fetal bovine serum (FBS)+10% dimethyl sulfoxide (DMSO). Once thawed, the cells were centrifuged at 200× g for 10 min and cultured in 24 well-plates in StemPro-34 complete medium (Thermo Fisher Scientific) supplemented with 200 mM GlutaMAX (Thermo Fisher Scientific), 1% penicillin/streptomycin (Thermo Fisher Scientific), 100 ng/mL Stem Cell Factor (SCF, Prepotech), 100 ng/mL FLT-3 (Thermo Fisher Scientific), 20 ng/mL Interleukin-6 (IL-6) (Thermo Fisher Scientific), 20 ng/mL Interleukin-3 (IL-3) (Peprotech). The medium was changed daily. On day 4, integration-free vectors encoding for the Yamanaka’s genes (CytoTune-iPS 2.0 Sendai Reprogramming Kit, Thermo Fisher Scientific) were used to transduce PBMCs. 14-21 days after, optimal hiPSC-like colonies were identified, picked and transferred onto Geltrex-coated wells. Cell maintenance was performed in feeder-free conditions with complete Essential (E8; Thermo Fisher Scientific). The spent medium was daily refreshed and passaging was performed at ∼ 85% confluency via EDTA-based dissociation solution.

### Embryoid Bodies Differentiation Assay

hiPSC clumps were plated on Pluronic acid-treated dishes in E8 medium supplemented with 5μM Y27632 ROCK inhibitor. After 48 h, the medium was replaced with a 1:1 mix of Essential 6 Medium (E6) (Thermo Fisher Scientific) and E8 medium. On day 4, cultures were shifted to E6 medium, and the medium was replaced daily. On day 7, EBs were collected and transferred on Geltrex-coated plastic, allowing them to adhere and spontaneously differentiate for additional 2 weeks. Over this period, the medium was replaced every other day and progressively substituted with DMEM High Glucose medium supplemented with 10% FBS (Sigma-Aldrich). On day 21, EBs were fixed or lysed for further analysis.

### RNA isolation and quantitative-PCR (qPCR)

Total RNA was isolated with Trizol Reagent (Thermo Fisher Scientific). 1μg of RNA was employed as template for reverse transcription (iScript cDNA Synthesis Kit, BioRad) according to manufacturers’ instructions. RT-qPCR was carried out on a CFX96 Real Time Thermalcycler (BioRad) via SsoAdvanced Universal SYBR Green Supermix Kit. Data were analyzed with a BioRad CFX Manager dedicated software and normalized using GAPDH house-keeping gene.

### Immunofluoresce Assay

HiPSCs and EBs were fixed with 4% PFA for 15 min RT, permeabilized with 0.5% Triton X-100 in PBS for 15 min RT and blocked with 5% FBS + 0.3% Triton X-100 in PBS for 1h RT. Samples were incubated with primary antibodies at 4°C overnight and then the signal was revealed with appropriate AlexaFluor secondary antibodies (primary and secondary antibodies used in this study are reported in Supplementary Table 1). Nuclei were counterstained with Hoechst 33258 1 μg/ml (Thermo Fisher Scientific). Cultures were inspected with the microscope Leica DM IL Led Fluo and pictures acquired with camera Leica DFC450 C (Leica Microsystem).

### Karyotyping and a-CGH of hiPSC clones

The cytogenetic analysis was performed using QFQ-banding techniques on metaphase chromosomes obtained by standard procedures from peripheral blood lymphocytes and from iPSC clones. At least 30 metaphases were scored in each case. Blood and iPSC genomic DNA were extracted through Promega Wizard™ Genomic DNA Purification Kits (Promega, Madison, WI, USA). Array comparative genome hybridisation (array-CGH) was performed on blood and hiPSC genomic DNA using the SurePrint G3 Human CGH Microarray Kit 4 x 180K in accordance with the manufacturer’s instructions (Agilent Technologies, Palo Alto, CA, USA). Probe positions are referred to human genome assembly GRCh37/hg19.

## RESULTS

In this study we selected a cohort of patients with a spectrum of clinical SCZ phenotypes, summarized in Table 1.

**Table 1.**
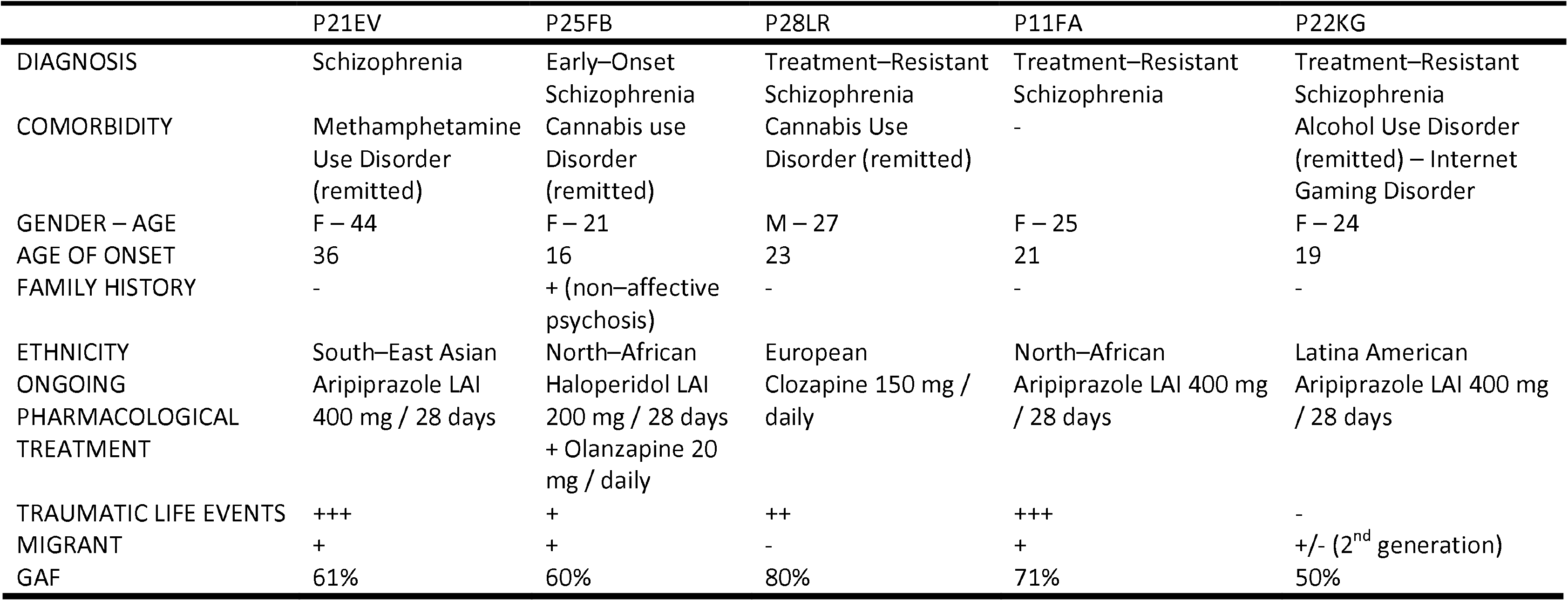
Summary of different clinical phenotypes in the SCZ study population.

From these SCZ patients, we isolated peripheral blood mononuclear cells (PBMCs) and transduced with non-integrating Sendai virus particles delivering the four Yamanaka reprogramming factors (OCT3/4, SOX2, c-MYC and KLF4) as previously reported (Marcatili et al; 2016, Marsoner et al; 2016). 10-12 days post-SeV transduction, clones appeared in culture. One week later, colonies were picked, expanded and their morphology monitored over time. Several clones from each reprogrammed PBMC sample were obtained and, for each subject, one clone was fully characterized. All generated clones gave rise to stable expanding hiPSC lines with characteristic human embryonic stem cell-like morphology, high nucleus/cytoplasmic ratio and colonies exhibiting defined borders (Figure 1A). Patient-derived iPSC characterization and validation is reported in Supplementary Table 2. To evaluate the pluripotent state of the hiPSC clones, immunofluorescence, and quantitative RT-PCR assays were performed. P28LR, P25FB, P21EV, P11FA, and P22KG SCZ-iPSC lines (Figure 1 B), as well as all the other generated lines (Supplementary Figure 1), showed immunoreactive signal for nuclear (NANOG, OCT4, SOX2) and surface (TRA-1-60) endogenous pluripotency markers. Transcripts for pluripotency-associated markers were also detected via RT-qPCR assay (Figure 1C). To rule out the impact of possible microbial contaminations, hiPSC cultures were tested for the absence of mycoplasma and all the lines were found negative. Finally, the functional pluripotency of the hiPSC clones was assessed by means of embryoid body (EB) formation assay. All patient-derived hiPSC lines, cultured for 7 days in suspension followed by 14 days in adhesion condition, underwent a transition from pluripotent stem cells into a heterogeneous population of cell types characterized by a variety of different morphologies. Notably, the cultures presented immunoreactive cells for endodermal (GATA 4), mesodermal (α-SMA), and neuroectodermal (TUBB3, also known as βIII-Tubulin) derivatives as detected by immunofluorescence staining (Figure 2A, Supplementary Figure 2). The subsequent gene expression analysis revealed substantial expression levels of transcripts for the above–mentioned markers belonging to the three germ-layers (Figure 2B).

**Figure 1.**
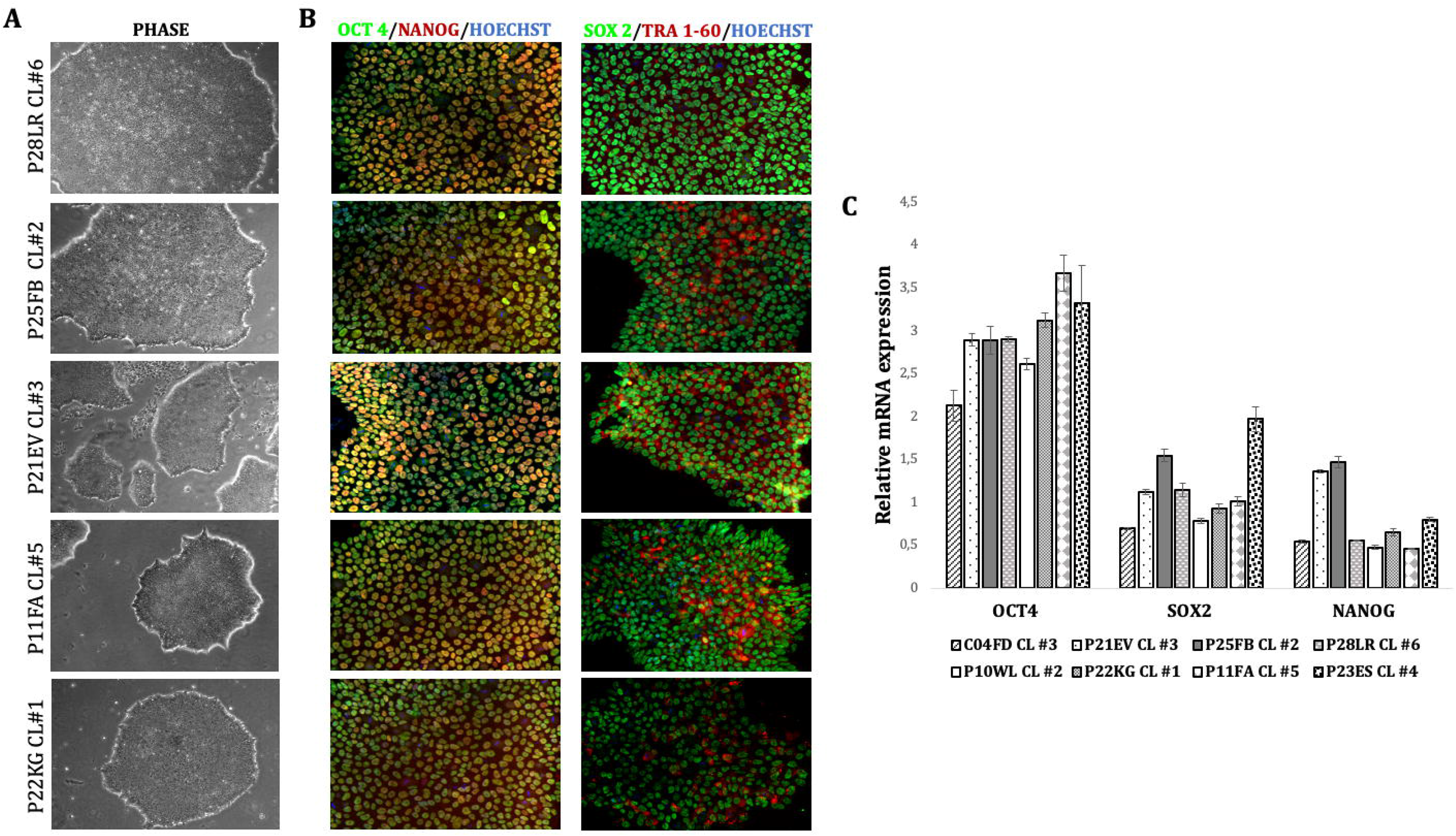
Characterization of SCZ-hiPSC clones. (A) Representative phase microscopy picture of patient-derived hiPSC colonies (5X). (B) Immunofluorescence showing the correct protein nuclear expression of OCT4 (green), NANOG (red), SOX2 (green) and surface localization of TRA-1-60 (red) pluripotency markers (20X). (C) RT-qPCR evaluating pluripotency-associated gene expression levels. Data were normalized on the house-keeping gene *GAPDH*.

**Figure 2.**
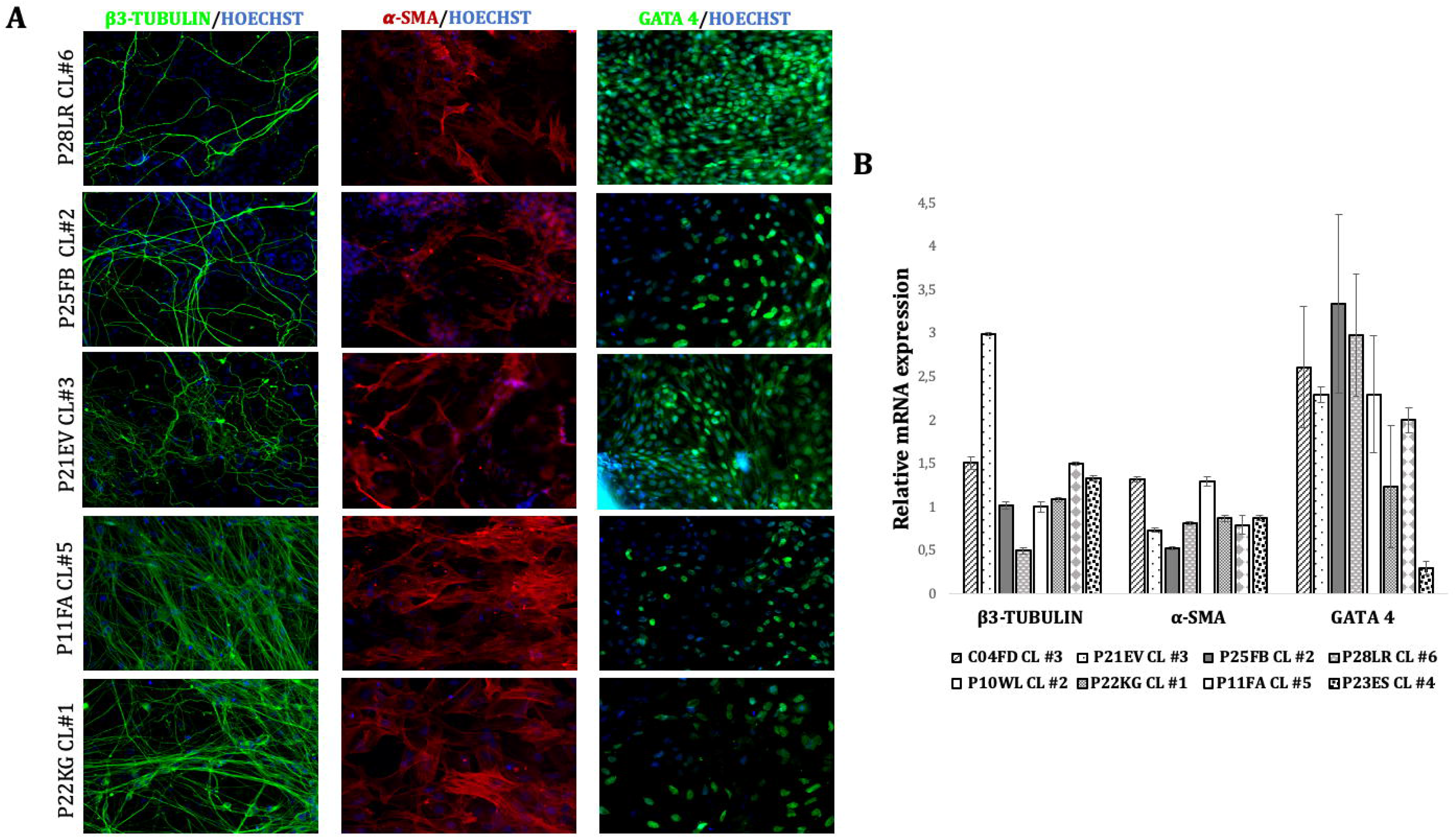
*In vitro* EB differentiation assay for patient-derived iPSC lines. (A) Differentiated 21-day old embryoid body cultures exhibit cells immunoreactive for neuro-ectodermal (β3-Tubulin), mesodermal (α-SMA) and endodermal (GATA4) lineage markers (20X). (B) RT-qPCR analysis displaying comparable genes expression levels among the different cultures. Data were normalized on the house-keeping gene *GAPDH*.

### Genetic characterization of hiPSC clones

One hiPSC clone was characterized for each individual patient by karyotyping and array CGH. Compared to patients’ PBMCs, no chromosome variation was found for the analysed hiPSC clones.

## DISCUSSION

Here we report the generation of a collection of hiPSC lines from SCZ patients with different clinical profiles, obtained through a stepwise reprogramming approach from PBMCs. Three TRS patients had different response profiles to clozapine, the only evidence-based pharmacological option for this condition, which affects around 30% of SCZ patients (Kane and Correll, 2016). Among several peculiarities, clozapine modulates glutamatergic transmission, has low affinity for dopamine D2 receptors, a relatively high affinity for D1 and D4 receptors compared to other antipsychotics, and has been shown to influence transcription and synaptic architecture (de Bartolomeis et al., 2022). In this context, patient-specific neural cells derived from iPSCs may help to clarify mechanisms that could inform the development of novel therapeutic strategies for TRS. One of our TRS patients was also diagnosed with comorbid gaming disorder; although no study systematically assessed the prevalence or incidence of this condition in patients with SCZ, several case reports suggest a potential link between excessive video gameplay or abrupt gaming disruption and psychosis in some patients, emphasizing the need for further research in this underexplored area (Huot-Lavoie et al., 2023).

Age of onset is also diverse in our collection. Whereas three patients had a typical early adulthood onset, one patient was diagnosed with EOS which is a rare, severe, and chronic form linked to premorbid developmental issues. It often shows poor response to neuroleptic treatment, higher hospitalization rates, and a challenging prognosis (Driver et al., 2020). On the other hand, one female patient had a relatively late, postpartum onset in her late thirties. Indeed, women diagnosed with SCZ may have a later onset of symptoms and better premorbid adjustment and coping strategies than men (Bucci et al., 2023).

Another strength of our study is the ethnic diversity of our study population, which reflects real–world practice. Available findings in SCZ research remain heavily biased toward Western, Educated, Industrialized, Rich, and Democratic (WEIRD) societies, restricting the applicability to diverse populations. This limitation carries implications for both clinical trials and genomic prediction models (Burkhard et al., 2021). Given the complex interplay of constitutive biology and environment in the pathogenesis of SCZ, identification of the different individual stressors experienced in higher and lower-income countries may contribute to specify their impact on predisposing biomolecular factors. Here, we provide specific hiPSC lines from individuals with variable ancestry, perceived childhood adversities, migrant and socioeconomic status, and substance misuse, all of which are well known to influence the course of SCZ (Jauhar et al., 2022).

HiPSC studies provide evidence of disrupted cortical development which begins during neurogenesis in SCZ. Differentiation of clinical profiles, treatment response and disease course is likely to shed light on the mechanisms which contribute to specific pathways, and perhaps support the development of individually tailored treatments.

## ACKNOWLEDGMENTS AND DISCLOSURES

This work was funded by the Italian Ministry of Health “Young Researchers – Theory Enhancing” grant (GR-2018-12367290) to AD’A and MO. AD’A, MO and LC conceived the study. MO and LC designed experiments, which were performed by EV, CVA and MMAS. EV analyzed data and prepared figures. LC and MO assessed the quality of data. LM, FL and CS collected the blood samples and wrote the first outline of patient histories. DB isolated and stored PBMCs. CG prepared karyotypes and PF ran CGH-array analyses. AD’A aligned the clinical and biomolecular information. AD’A, LC and EV wrote the manuscript. We thank Anna Caretti for full access to laboratory equipment at the Department of Health Sciences. The results of this study were partially presented at the 13^th^ Congress of the Italian Society for Biological Psychiatry in 2023. The authors report no biomedical financial interests or potential conflicts of interest.

ARTICLE INFORMATION From the Department of Mental Health and Addiction, ASST Santi Paolo e Carlo, Milan, Italy (EV, AD’A); Department of Cellular, Computational, and Integrative Biology (CIBIO), University of Trento, Italy (EV, CVO, LC); Departments of Health Sciences (DB, CG, LM, FL, CS, AD’A) and of Medical Biotechnology and Translational Medicine (PF), University of Milan, Italy; Department of Biology, Unit of Cell and Developmental Biology, University of Pisa, Italy (MMAS, MO).

---

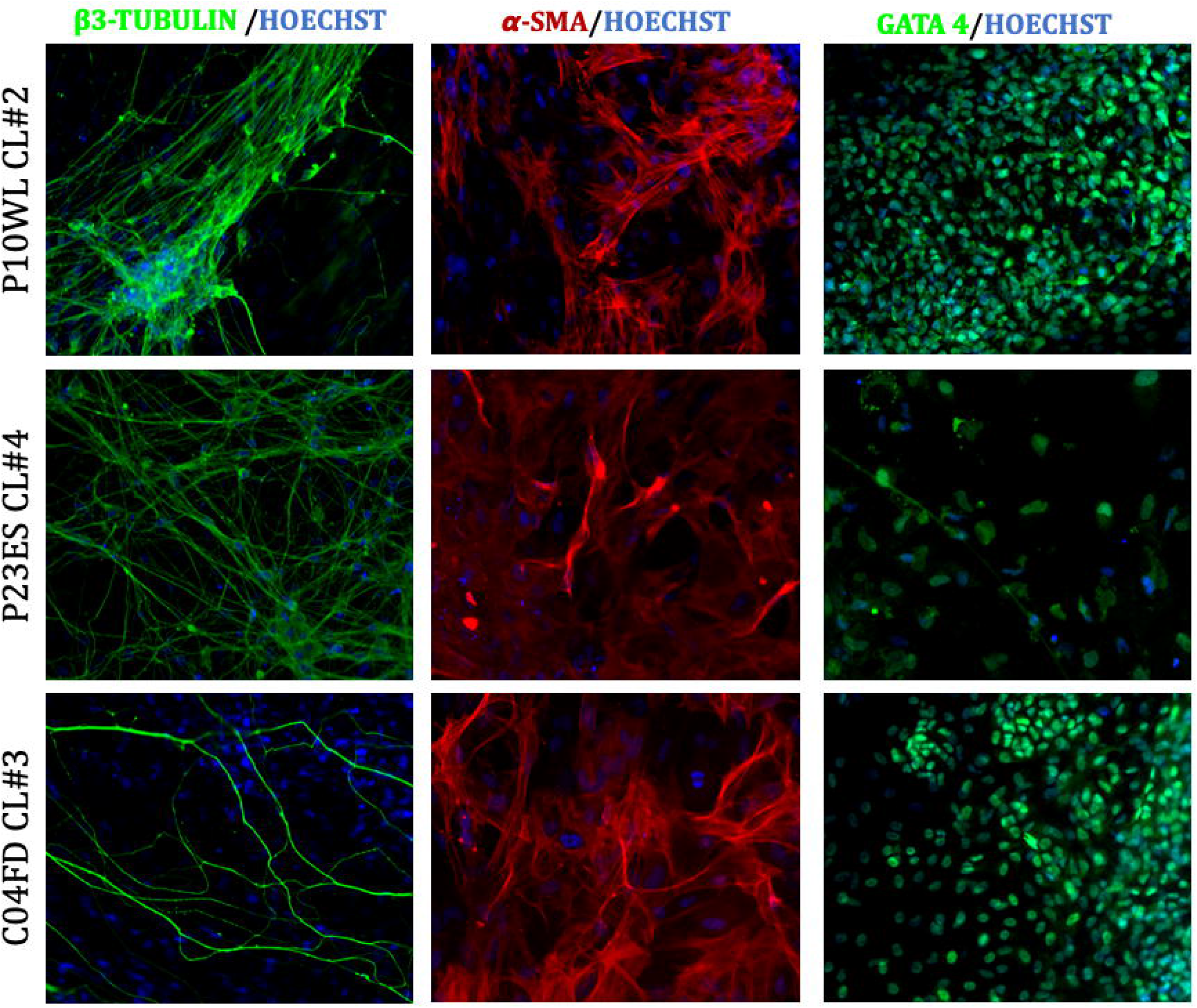

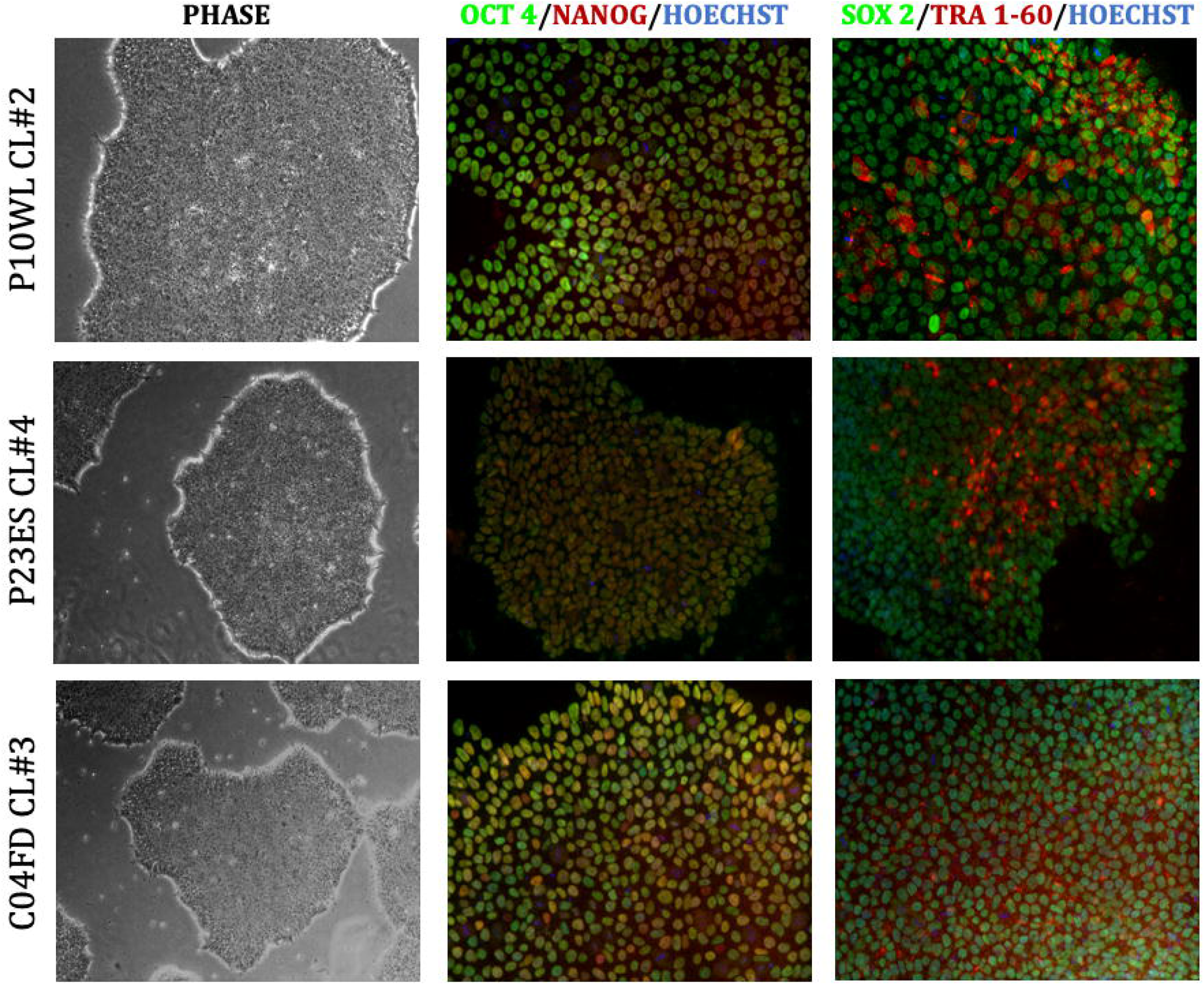

